# AttentionPert: Accurately Modeling Multiplexed Genetic Perturbations with Multi-scale Effects

**DOI:** 10.1101/2024.02.02.578656

**Authors:** Ding Bai, Caleb Ellington, Shentong Mo, Le Song, Eric Xing

## Abstract

Genetic perturbations (i.e. knockouts, variants) have laid the foundation for our understanding of many diseases, implicating pathogenic mechanisms and indicating therapeutic targets. However, experimental assays are fundamentally limited in the number of perturbation conditions they can measure. Computational methods can fill this gap by predicting perturbation effects under unseen conditions, but accurately predicting the transcriptional responses of cells to unseen perturbations remains a significant challenge. We address this by developing a novel attention-based neural network, AttentionPert, which accurately predicts gene expression under multiplexed perturbations and generalizes to unseen conditions. AttentionPert integrates global and local effects in a multi-scale model, representing both the non-uniform system-wide impact of the genetic perturbation and the localized disturbance in a network of gene-gene similarities, enhancing its ability to predict nuanced transcriptional responses to both single and multi-gene perturbations. In comprehensive experiments, AttentionPert demonstrates superior performance across multiple datasets outperforming the state-of-theart method in predicting differential gene expressions and revealing novel gene regulations. AttentionPert marks a significant improvement over current methods, particularly in handling the diversity of gene perturbations and in predicting out-of-distribution scenarios.

## 1 Introduction

Single-cell RNA-sequencing (scRNA-seq) has advanced significantly, enabling the generation of transcriptomic datasets encompassing millions of cell samples [28, 1, 8, 15]. This advancement facilitates CRISPR-based perturbational screens and rapid experimental sampling of genetic perturbation outcomes [7, 22, 11, 24]. However, this field faces significant challenges, particularly when expanding the number of genes for perturbation. The potential combinations of two-gene perturbations grow quadratically with an increasing number of genes [22], leading to a combinatorial explosion in scenarios involving even more simultaneous gene perturbations. Consequently, these methods are constrained by scalability and cost-effectiveness, especially when exploring new genetic perturbations [7, 6, 22].

Many computational simulations model the gene regulatory networks (GRN), rather than directly incorporating the effects of perturbations, ranging from traditional Bayesian networks [12] and causal inference algorithms [32] to more contemporary models such as SENIC [2] and CellOracle [17]. Recent studies [23, 27] have indicated that achieving accurate GRN inference from transcriptomic data poses significant challenges, and GRN-predicting models often fall short in performance when tasked with predicting outcomes of single-gene and multi-gene perturbations. Moreover, other mechanistic modeling methods [13, 35] and linear models [7, 17] addressing both chemical and genetic perturbations, face limitations including scalability issues in predicting responses for a vast number of genes, or an inability to capture non-linear dynamics.

Current advancements in the field use deep neural networks trained on datasets encompassing extensive cells and genes with a moderate number of perturbations, which map genes, perturbations, or genetic relationships into a latent space for the prediction of perturbation outcomes [21, 20, 10, 25, 29, 27, 34]. Most of these methods, including scGEN [21], PerturbNet [34] and CPA [20], primarily predict the transcriptional effects of chemical treatments rather than genetic perturbations. They often approach genetic perturbations as analogous to varying doses, which inadequately capture the complex interactions between different genes and the perturbed ones, resulting in limited performances in this particular domain [20].

Recently, there has been a growing trend in employing large language models (LLMs) trained on unlabeled single-cell transcriptomic datasets, which often takes predicting perturbational effects as one of their downstream tasks [14, 16, 5, 33]. While these LLMs represent a relative advancement over other methodologies, they also benefit from downstream methods by incorporating expression representations generated by the foundational LLMs as inputs. Furthermore, the training of these foundation models requires extensive resources, utilizing large-scale datasets and numerous GPUs, which incurs significant time and financial costs.

Among all computational methods for genetic perturbation effects prediction, the state-of-theart (SOTA) method graph-enhanced gene activation and repression simulator (GEARS) [27], in contrast, specifically targets genetic perturbations. GEARS significantly surpasses other methods in this area, illustrating the effectiveness of a genetic-perturbation-centric preprocessing and training approach for future methodologies [27]. Current single-cell LLMs often take GEARS as a downstream probe in predicting perturbational effects [14, 16, 5, 33]. A key innovation of GEARS is its incorporation of a highly expressive and informative knowledge graph based on Gene Ontology similarities. Nonetheless, GEARS as well as other methods still exhibit the three primary limitations that our work aims to address and advance.

We propose **AttentionPert**, a multi-level deep-learning method to accurately predict transcriptional responses to multiplexed genetic perturbations, which integrates the multi-head attention mechanism [3] with graph neural networks on augmented gene interactions, alongside pretrained co-expressive gene representations. AttentionPert is proposed to improve three major detriments of current deep-learning works on this problem.

- AttentionPert primarily improves existing methods by capturing uneven effects that the perturbation in a particular gene has on various other genes. The attention-based **PertWeight** encoder perturbs all the gene-representing vectors in high-dimensional latent space with non-uniformly weighted offsets. PertWeight is designed to learn both the overarching background and distinct gene-specific global perturbational effects, reflecting the complex gene-gene interactions within the gene regulatory network.
- AttentionPert accounts for localized gene perturbation analysis using the **PertLocal** encoder, complementing the global effects learned by PertWeight. PertLocal is designed to learn the local disturbance in the representation space of perturbed genes on themselves and their neighbors in a knowledge graph of genetic regulations and interactions. The PertLocal encoder also avoids a pure linear combination of effects of perturbed genes and hence learns the non-additive coeffects of multi-gene perturbations.
- AttentionPert utilizes a pre-trained context gene representation to initialize all the gene embeddings rather than random initialization. This approach significantly improves its capacity to predict responses to unseen perturbed genes.

AttentionPert outperforms prior methods including the SOTA GEARS model on *RPE1* and *K562* datasets [26], each of which comprises more than 1, 000 single-gene perturbations, and on the *Norman* dataset containing 131 two-gene perturbations [22]. Particularly, AttentionPert has exceptional capability in the most challenging out-of-distribution (OOD) task: predicting the effects of novel multi-gene perturbations where none of the perturbed genes have been encountered during the training phase. This remarkable performance highlights its ability to generalize effectively to entirely new genetic scenarios. Our experiments also reveal insights into nuanced genetic interactions associated with multi-gene perturbations through detailed gene-specific and perturbation-specific analyses.

## 2 Method

### 2.1 Problem setting

AttentionPert predicts post-perturbation gene expressions given specific gene-perturbation conditions. Formally, for a gene-perturbation condition denoted as *c* = *{j*_1_, *j*_2_, …, *j*_*m*_*}*⊆ *{*1, …, *K}*, representing a subset of gene indices, the objective is to predict the post-perturbation gene expression distribution *Y*_*c*_ ∈ ℝ^*K*^. Within the datasets, each perturbation is associated with multiple cells, of which post-perturbation expressions are represented as 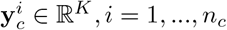, where *n*_*c*_ is the number of cells corresponding to the condition *c*. A perturbation condition is termed *control* when *m* = 0 and the unperturbed expressions are considered as known entities denoted by *{***x**^*i*^, *i* = 1, …, *n*_control_*}*. The primary goal for a parametric model parameters *Y*_*θ*_ is to minimize the difference between the expected *Y*_*θ*_(*c*) and the mean of the ground truth samples 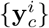 for each perturbation *c*.

Existing deep neural network models process both the perturbation condition and a randomly sampled unperturbed cell to predict a post-perturbation expression vector [20, 27]. These predicted vectors are evaluated by measuring their similarity to actual perturbed samples. In our method, we adhere to the established preprocessing stage by randomly pairing each control vector 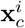 from the set {**x**^*t*^, *t* = 1, …, *n*_control_} with its corresponding post-perturbation vector 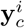, where *i* = 1, …, *n*_*c*_ for each condition *c*. While CPA encodes gene expressions into a high-dimensional space and then perturbs them within this space [20], our approach follows GEARS, focusing on directly predicting the expression shift resulting from the perturbation condition and adding it to the control expression, formally expressed as: 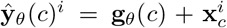. We also adopt the combined loss function used by GEARS, which integrates auto-focus and direction-aware loss, to compare our predictions ŷ_*θ*_(*c*)^*i*^ with the ground-truth 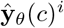. We detail the preprocessing and loss function formulas in Supplementary Section A.1. The primary innovation of our model is the improved post-perturbation expression change predictor **g**_*θ*_.

### 2.2 Model overview

AttentionPert takes a perturbation condition *c* ⊆ *{*1, …, *K}* as input. The vector **z**_*c*_ ∈ *{*0, 1*}*^*K*^ contains the 0/1 perturbation indicators, formally: **z**_*c,j*_ = 1 if *j* ∈ *c* and 0 otherwise. AttentionPert feeds the perturbation indicators into two modules called PertWeight and PertLocal encoders respectively. Each encoder outputs a matrix of encoding, noted as **H**^PW^ by PertWeight and **H**^PL^ by PertLocal, which both ∈ ℝ^*K×D*^, where *D* is the network hidden size. After that, a genespecific decoder takes the sum of two encodings and then generates the predicted post-perturbation expression shift **g**_*θ*_(*c*), as shown in Figure 1. Finally, we add the shift to a random control sample 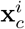 to get a post-perturbation prediction 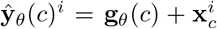, and compute the loss function for the whole batch to learn the model parameter *θ*. A more intuitive illustration of this method is Supplementary Section A.2.

**Figure 1.**
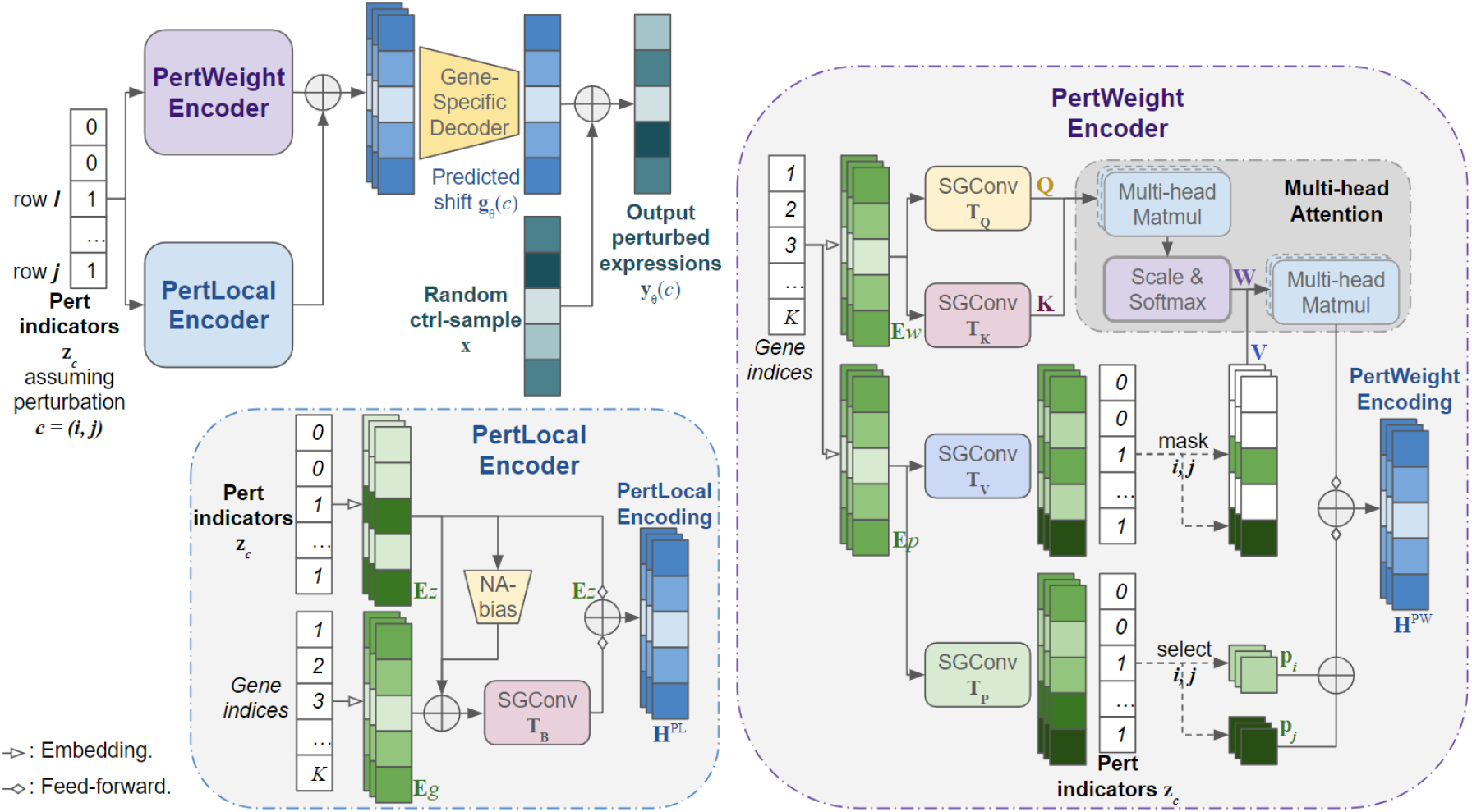
The overall architecture of AttentionPert. The illustration assumes that the perturbation condition *c* = *{i, j }*. The indicators vector **z**_*c*_ is the input of both the PertWeight Encoder and the PertLocal Encoder. PertWeight generates attention weight matrices **W** based on the embedding **E**_*w*_, uses the perturbation indicators to filter the values **V** and **P**, and then combines both the weighted and the general global effects. PertLocal takes the perturbation indicators and outputs the locally disturbed gene states. The gene-specific linear decoder decodes the sum of two encodings into the perturbation effects, and then the predicted expressions are obtained. All SGConvs of PertWeight: **T**_*Q*_, **T**_*K*_, **T**_*V*_ and **T**_*P*_ take the Gene Ontology graph as the input graph, while the SGConv of PertLocal **T**_*B*_ takes the augmented GO graph. All embeddings for gene indexes are initialized using pre-trained Gene2Vec embeddings and can be learned during training. All the feed-forward layers are multiple linear and batch normalization layers.

### 2.3 PertWeight for non-uniform global effects

Current works treat all affected genes uniformly when computing the effects of a genetic perturbation in the embedding space, which does not differentiate how a perturbed gene might impact other genes in unique ways [21, 20, 27]. To address this issue, the attention-based perturbation encoder **PertWeight** is designed to learn weight matrices representing the interactions from perturbed genes to other genes. These matrices are then applied in a weighted addition to compute post-perturbation states, thus enabling the model to capture the non-uniform gene-to-gene perturbational effects.

PertWeight firstly embeds gene indices *{*1, 2, …*K}* into 2 independent sets of embeddings. We define the first embedding as the *weight* embedding noted by **E**_*w*_, and the second embedding as the *pert* embedding noted by **E**_*p*_. Here: **E**_*w*_, **E**_*p*_ ∈ ℝ^*K×D*^*e*. Both embeddings can be either randomly initialized or loaded with the pre-trained gene-representing embeddings. Current studies assign each gene a high-dimensional representation that is randomly initialized [20, 27], which is a suboptimal approach considering the limited range of experimental perturbation conditions available in comparison to the potential perturbations. Utilizing pre-trained context representations for initializing gene embeddings could significantly enhance the model’s ability to predict responses to unseen perturbed genes. Our proposed method incorporates Gene2Vec [9], which offers 200-dimensional vector representations for human genes, derived from gene co-expression patterns across 984 datasets in the Gene Expression Omnibus (GEO) databases. It is proven to be more expressive through our experiments than random initializing gene representations.

In this module, we apply 4 convolutional graph neural network (GNN) encoders with the Gene Ontology (GO) graph as the input graph 𝒥 with an adjacency matrix **J**. The GNN model *SGConv* [19] has linear projection weights 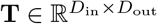. Taking the input graph node embedding 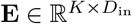 and the graph 𝒜 of *K* nodes with an adjacency matrix **A** ∈ ℝ^*K×K*^, a *t*-hop SGConv outputs a graph-convolution encoding 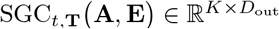.

A Gene Ontology term provides a structured framework to represent information in a specific domain, where every gene is associated with multiple Gene Ontology terms [4, 31, 30]. The GO graph 𝒥 is constructed through Gene Ontology following the SOTA method [27], where the edge weight of two genes is the fraction of their shared Gene Ontology terms. Details for the GO graph are shown in Supplementary Section A.3

The 4 SGConv encoders have different weight matrices: 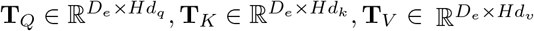 and 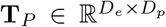 respectively, where *H* is the number of heads in the multi-head attention, and *d*_*q*_ = *d*_*k*_. SGConvs **T**_*Q*_ and **T**_*K*_ process the weight embedding **E**_*w*_ and SGConvs **T**_*V*_ and **T**_*P*_ process the pert embedding **E**_*p*_. Then we got 4 outputs:

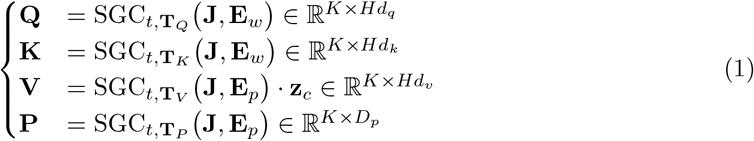

Note that here the output of the SGConv **T**_*V*_ is masked using the vector of perturbation indicators **z**_*c*_, leaving only the values of perturbed genes. **Q, K** and **V** are then fed into the multihead attention as queries, keys and values [3], as shown in Figure 1. Specifically, queries, keys and values are all equally divided into *H* matrices along the second dimension. Note that for head *h*, the sub-matrices are **Q**^*h*^, **K**^*h*^ and **V**^*h*^, and the attention matrix of each head is:

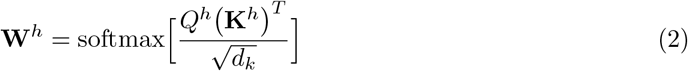

And the output of the multi-head attention **H**^MHA^ is the concatenation of all heads **H**^MHA,*h*^ = **W**^*h*^**V**^*h*^. Since **V** is masked by the vector of perturbation indicators **z**_*c*_, the output can also be written as:

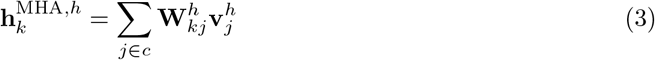

for each gene *k* ∈ *{*1, …, *K}*. This is a weighted summation of perturbational effects from all perturbed genes. All attention matrices **W**^*h*^ ∈ **R**^*K×K*^ are the *Perturbation Weight Matrices*, which are the key innovation of PertWeight module featuring the various effects between any perturbed genes and all genes.

The output **P** represents the general effects of perturbed genes upon all genes. The row vectors of **P** in the perturbed genes *{***p**_*j*_ |*j* ∈ *c}* are selected and added to the final output. Hence the final output of the PertWeight Encoder is:

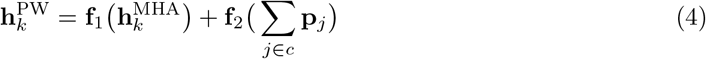

for each gene *k*, where **f**_1_ and **f**_2_ are linear feed-forward layers that project encodings into the same hidden size *D*, followed by layer-normalization. The output matrix is 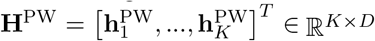.

Obviously, PertWeight is additive for genes in any multi-gene perturbation *c*, which cannot simulate the non-additive feature, of which details are shown in Supplementary Section A.4. Therefore, we propose PertLocal which learns the nonlinear co-effects of perturbations of multiple genes.

### 2.4 PertLocal for local disturbance

Many existing models primarily focus on the perturbational effects across all genes within a cell [21, 20], overlooking the different interactions between genes. The SOTA method employs a genegene interaction graph, yet it only models global effects [27]. However, genetic regulation studies demonstrate that local and global perturbational effects complement each other. **PertLocal** is designed to overcome this gap by learning the localized disturbance effects of perturbed genes on their closely related counterparts. It complements the global effects captured by the PertWeight encoder, thereby significantly enhancing the overall predictive accuracy of the model.

Similar to PertWeight, PertLocal also starts with 2 embeddings. The first one is the gene base embedding representing unperturbed states of genes that embeds gene indices *{*1, 2, …*K}* into dimension *D*_*e*_, and the second one is the perturbation indicator embedding that embeds binary indicators *{*0, 1*}* into dimension *D*_*e*_. The indicator embedding layer embeds the perturbation indicator vector **z**_*c*_ ∈ *{*0, 1*}*^*K*^ into a matrix of dimension *K × D*_*e*_. We denote the gene base embedding as 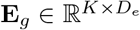 and the perturbation indicator embedding as 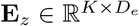. Similarly to **E**_*w*_ and **E**_*p*_, the gene base embedding **E**_*g*_ is also initialized with Gene2Vec.

Numerous previous studies have inaccurately employed the additive prediction approach commonly used in multi-gene perturbation modeling [12, 32, 2, 17], which the SOTA work designed an innovative cross-gene layer to avoid [27]. In contrast, PertLocal adopts a non-additive bias layer to effectively learn the complex, non-linear coeffect resulting from multiple perturbed genes. The non-additive (NA) biased embedding for the perturbation indicator embedding is formally:

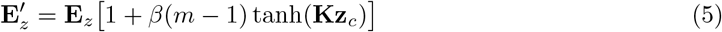

where *β* ∈ (0, 1) is a hyper-parameter, *m* = *c* is the number of perturbed genes, and 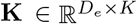 is a trainable weight matrix. PertLocal then applies another *t*-hop SGConv encoder on the summation of the 2 embeddings 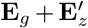, which takes the augmented Gene Ontology graph as the input graph. The augmented GO graph 𝒥^*′*^ is created by adding virtual edges to all disconnected pairs of nodes in the GO graph with a new minimum edge weight. The process to attain the GO graph and the augmented GO graph is shown in Figure 2. Formally,

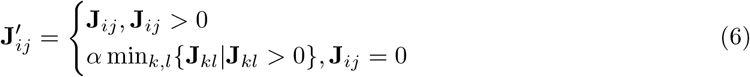

where 0 *< α <*= 1. Virtually added edges with minimum weights make the graph 𝒥^*′*^ fully connected, which allows PertLocal to learn lightweight instead of zero global gene-gene reactions. The details of the GO graph and the GNN encoders can be found in Supplementary Section 2.

**Figure 2.**
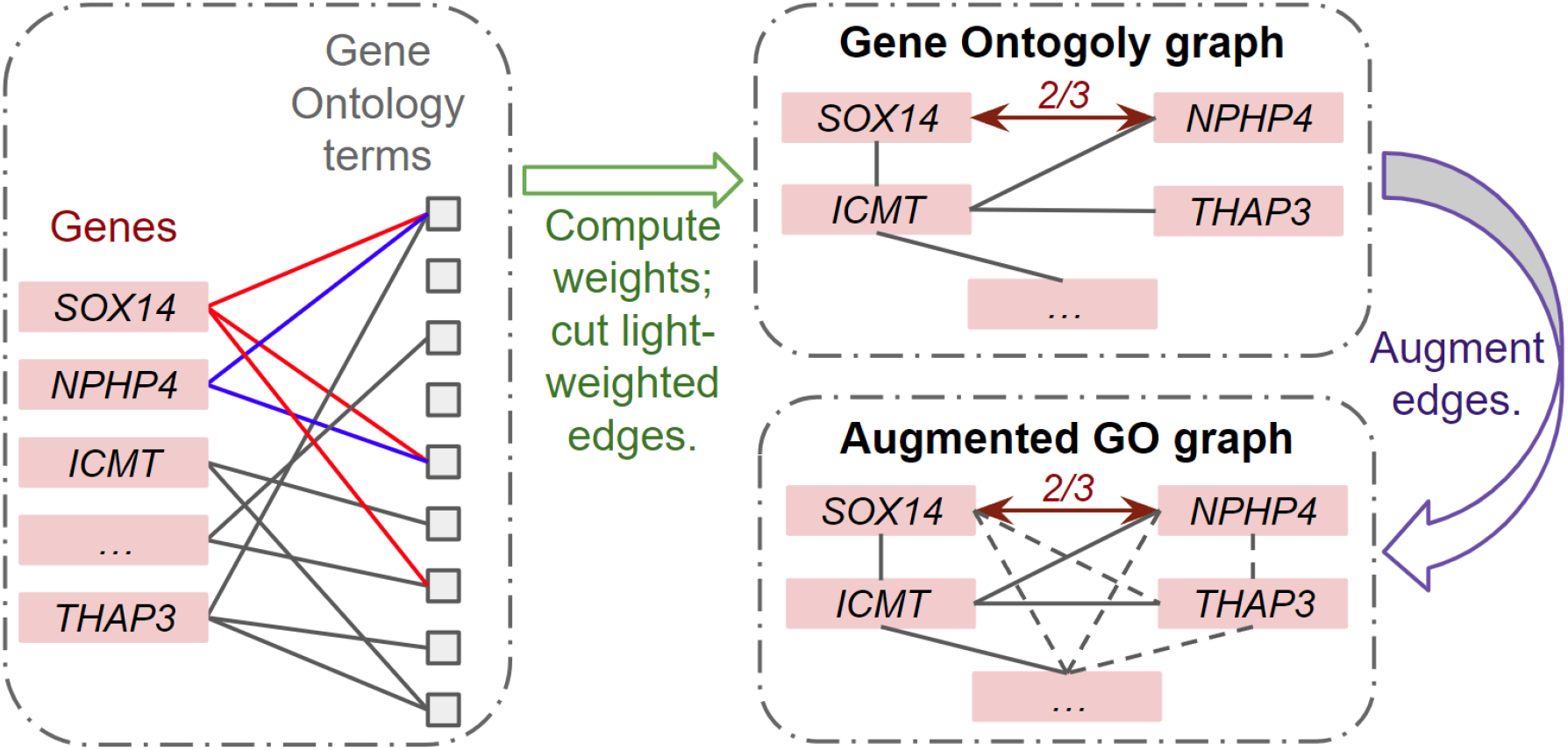
The process to obtain the Gene Ontology (GO) graph and the augmented GO graph. Using the GO terms of each pair of genes, for example, those marked in red for gene SOX14 and those marked in blue for NPHP4, the edge weight is defined as the number of shared GO terms divided by the size of the union set of their GO terms, which is 2*/*3 for genes SOX14 and NPHP4 (only as an illustration in the figure). The GO graph is then obtained by filtering 20 highest edge weights for each gene and cutting all other edges. The augmented GO graph is fully connected, in which each edge either exists in the GO graph or is augmented as *α* times the minimum edge weight of the GO graph, where *α* ∈ (0, 1].

Denote the weight matrix for the PertLocal SGConv as 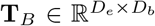. After 2 feed-forward and then batch normalization layers **f**_3_ and **f**_4_ of the same output size *D*, the SGConv output and the perturbation indicator embedding are then added together as the output of the PertLocal Encoder:

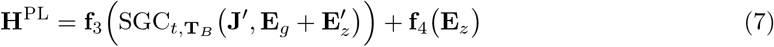

as shown in Figure 1.

The PertLocal encoder is non-additive since the NA-biased embedding is identical to the indicator embedding when *m* = 1 but not when *m* ≥ 2, which then induces the non-additive property (Supplementary Section 1.4 A.4). Through the weighted gene-gene graph structure and the indicator embedding, PertLocal learns the representations of local perturbational effects that complement the PertWeight encoding.

### 2.5 Decoder for transcriptional effects

After getting the encodings from both encoders, we input the sum of **H**^PW^ and **H**^PL^ into a linear decoder, which simplifies GEARS decoder by removing its cross-gene part. In this decoder, after a feed-forward function for each gene *k*, a gene-specific linear layer maps the post-perturbation encoding into a scalar that is the predicted perturbation effect. Formally:

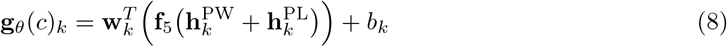

where **w**_*k*_ ∈ ℝ^*D*^ and *b*_*k*_ ∈ ℝ are learnable weights and bias of the linear projection for each gene. The cross-gene layer is used in GEARS for predicting the non-additive effects of multi-gene perturbations, which is covered in our PertLocal model. Since PertWeight and PertLocal have learned both the uneven global and the local perturbational effects, a relatively less expressive cross-gene layer is abandoned to avoid redundancy. Experiment results demonstrate that AttentionPert achieves superior results, outperforming the need for a cross-gene layer.

The final output is the predicted post-perturbation gene expressions ŷ_*θ*_(*c*)^*i*^ obtained by adding the predicted post-perturbation expression shift vector **g**_*θ*_(*c*) to a randomly sampled unperturbed control gene expressions 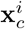, as shown in Figure 1. The combined loss function (inherited from GEARS) is computed by taking the outputs of a mini-batch and then optimized to learn the parameters of **g**_*θ*_.

## 3 Results

### 3.1 Experimental setup

#### Evaluation metrics

The evaluation metrics for each perturbation condition *c* is the mean squared error (MSE) between the mean prediction and the average ground-truth post-perturbation expression on the top 20 differentially expressed (DE) genes. We also evaluate the Pearson correlation coefficient between the predicted post-perturbation expression shift and the ground-truth shift *ρ*_Δ_. This metric is evaluated throughout all genes to test the model’s ability to predict the overall expression change. Detailed descriptions of metrics are shown in Supplementary Section B. **Baselines**

We conduct experiments for 3 baselines and compare their performances with AttentionPert. The first one is the model *Ctrl* that directly outputs the mean value of all unperturbed control expressions, which is hence the worst case for any dataset. The performance of *Ctrl* also indicates the difficulty of the post-perturbation prediction since it measures how much effect those perturbations have on unperturbed cells. Due to the differences between datasets and perturbations inside the same dataset, the difficulty of predicting the effect of novel gene perturbations varies much. Hence we also calculate the relative mean squared error (*rel-MSE*) for each model that is the MSE of the given model divided by the MSE of *Ctrl*. The other 2 baselines are CPA [20] and GEARS [27].

#### Implementation and training

AttentionPert is implemented with the embedding size *D*_*e*_ = 200 of all embeddings fit the Gene2Vec. All hyper-parameters of AttentionPert are set as (Supplementary Section C): the hop number of all SGConvs *t* = 1; the output dimension of SGConvs **T**_*P*_ and **T**_*B*_ equals to the network hidden size *D*_*p*_ = *D*_*B*_ = *D* = 64; the number of heads *H* = 2 and the head dimension *d*_*q*_ = *d*_*k*_ = *d*_*v*_ = 64 in the multi-head attention of the PertWeight encoder; the minimum edge weight of the GO graph is multiplied by *α* = 0.75 when augmenting it to the Augmented GO graph, and the NA-bias parameter *β* = 0.05. We train the model in each data split independently for 20 epochs, with a batch size 128 and an Adam optimizer with a learning rate 10^−3^. We run the training process for all models on 5 splits of 3 datasets that are Norman, RPE1 and K562 [22, 26], of which details are shown in Supplementary Section D. Except for *Ctrl* which is non-parameterized, we conduct each experiment 5 times with randomly different model initialization and take the average and standard deviation of the results.

### 3.2 Comparison to prior work

The high cost of CRISPR gene-perturbation screens requires an in-silico method that predicts transcriptomic effects of novel single/multi-gene perturbations not in the existing datasets. Therefore in our experiments, all the cells of a given perturbation condition *c* are either in the train and validation set or in the test set, which simulates the task of predicting unseen perturbations. For single-gene perturbations, we define only one generalization class called *seen 0/1* where the only perturbed gene is never seen in the train and validation set. When considering the combination of multiple perturbed genes, however, there are multiple testing scenarios, depending on the experimental exposure of the genes to perturbations during training.

AttentionPert is adept at predicting transcriptional outcomes for perturbations involving multiple genes, of which performance is evaluated using the Norman dataset [22] that comprised 131 two-gene perturbations. As previously mentioned, two-gene perturbations in the test set are divided into three generalization classes for evaluation purposes. The first class includes scenarios where both genes in a two-gene perturbation set are individually encountered in the training data (*2-gene perturbation, seen 2/2*). The more challenging scenarios involve cases where either one gene (*seen 1/2*) or neither gene (*seen 0/2*) of the two-gene set are perturbed in the training dataset. The comparative analysis of these three scenarios is detailed in Table 1. The results demonstrate that AttentionPert outperforms all baseline models across these scenarios, particularly in the most OOD scenario where both test genes are unseen in the train set.

**Table 1:**
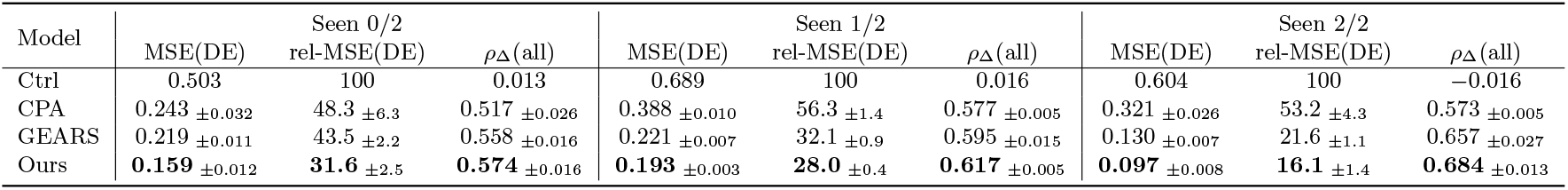
Seen 0/2, seen 1/2, and seen 2/2 comparison of 3 metrics on Norman dataset. MSE(DE) is the MSE of the top 20 DE genes. The relative MSE(DE) is in percentage (%) and is relative to the Ctrl model. The *ρ*_Δ_ score refers to the Pearson score between the predicted shift and the ground-truth shift. We record the mean and the standard deviation over 5 independent experiments. The best performances are marked in bold.

| Model | Seen 0/2 |  |  | Seen 1/2 |  |  | Seen 2/2 |  |  |
| --- | --- | --- | --- | --- | --- | --- | --- | --- | --- |
| | MSE(DE) | rel-MSE(DE) | $\rho_\Delta$(all) | MSE(DE) | rel-MSE(DE) | $\rho_\Delta$(all) | MSE(DE) | rel-MSE(DE) | $\rho_\Delta$(all) |
| Ctrl | 0.503 | 100 | 0.013 | 0.689 | 100 | 0.016 | 0.604 | 100 | -0.016 |
| CPA | 0.243 ± 0.032 | 48.3 ± 6.3 | 0.517 ± 0.026 | 0.388 ± 0.010 | 56.3 ± 1.4 | 0.577 ± 0.005 | 0.321 ± 0.026 | 53.2 ± 4.3 | 0.573 ± 0.005 |
| GEARS | 0.219 ± 0.011 | 43.5 ± 2.2 | 0.558 ± 0.016 | 0.221 ± 0.007 | 32.1 ± 0.9 | 0.595 ± 0.015 | 0.130 ± 0.007 | 21.6 ± 1.1 | 0.657 ± 0.027 |
| Ours | **0.159** ± 0.012 | **31.6** ± 2.5 | **0.574** ± 0.016 | **0.193** ± 0.003 | **28.0** ± 0.4 | **0.617** ± 0.005 | **0.097** ± 0.008 | **16.1** ± 1.4 | **0.684** ± 0.013 |

Furthermore, Figure 3 shows the specific comparison between AttentionPert and the SOTA method GEARS for each two-gene perturbation out of the 79 tested ones. Among them, AttentionPert outperforms GEARS on all 9 perturbations of the hardest seen 0/2 scenario. Over 3 scenarios, AttentionPert has better performances on 63 perturbations, which is about 80% of the test cases. This reveals that AttentionPert surpasses the SOTA method in predicting the effects of two-gene perturbations.

**Figure 3.**
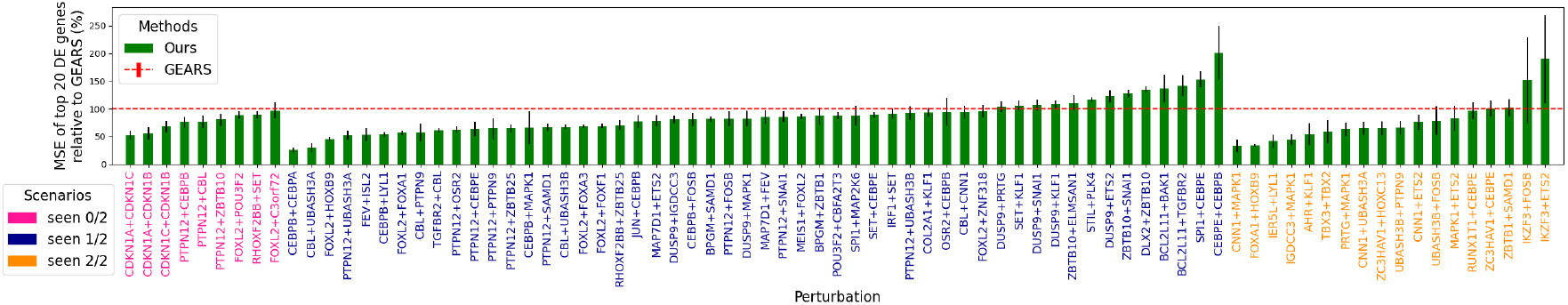
Perturbation-specific comparison on Norman dataset. The MSE(DE) values of AttentionPert are normalized as percentages relative to the performance of GEARS.

We apply two metrics evaluating the average error rate when dealing with different genes apart from MSE(DE) and *ρ*_Δ_(all) which represent overall performances across the set of genes. The first metric quantifies the proportion of the top 20 DE genes for a given perturbation predicted to change in an opposite direction compared to their actual change. The second metric is the percentage of DE genes for which the predicted post-perturbation gene expression falls outside one standard deviation of the actual post-perturbation expression distribution mean. Both metrics describe the ratio of wrongly predicted genes, of which lower values are preferable. Figure 4 shows the comparison between baseline models and AttentionPert on these two scores, demonstrating that our model makes fewer errors in predicting both the direction and value of perturbation effects for the top 20 DE genes.

**Figure 4.**
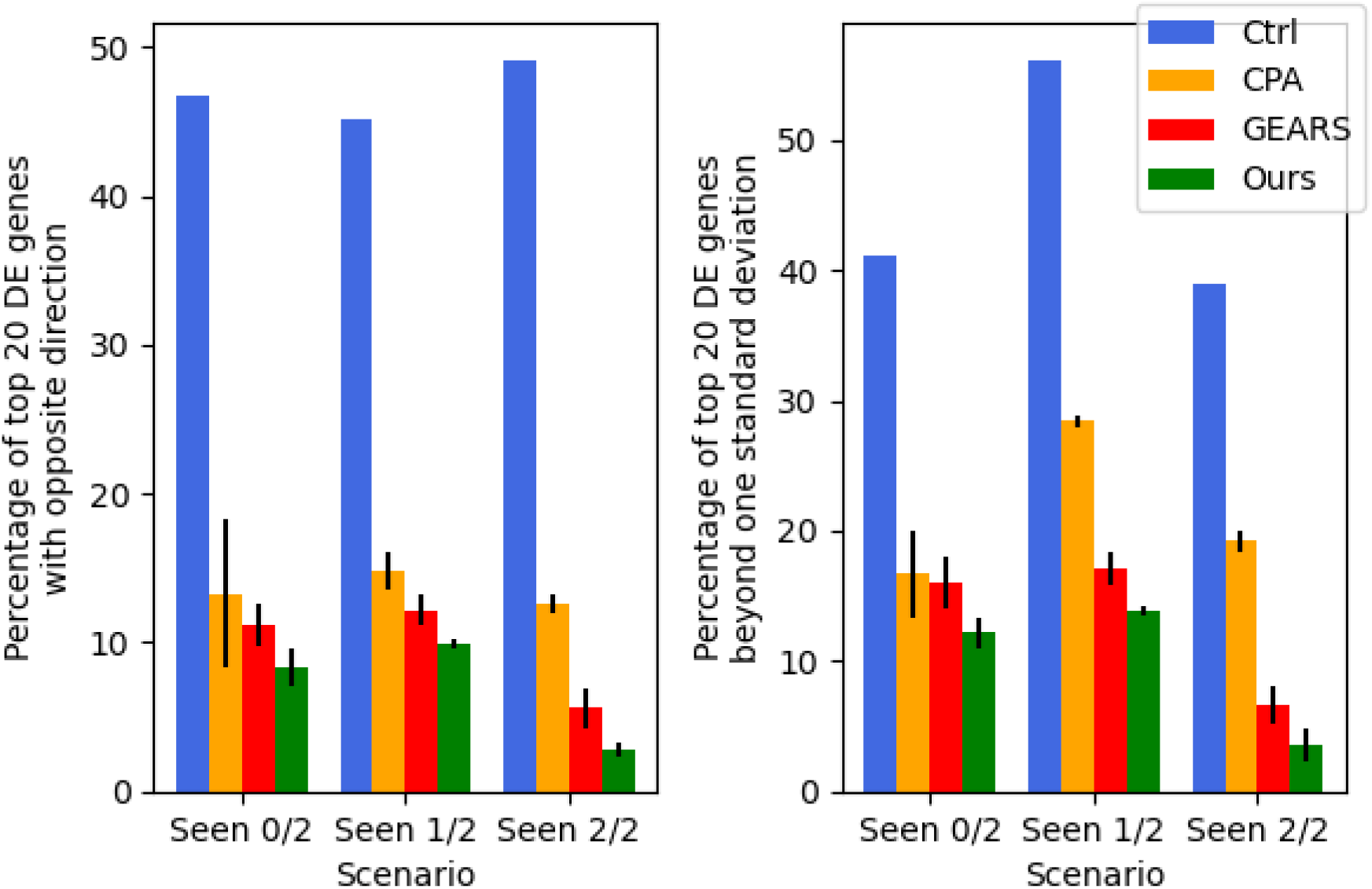
Comparison of metrics representing error rates on Norman dataset. These two metrics evaluate the average proportion of genes with incorrect direction and value predictions, respectively.

For single-gene perturbations, we employed datasets from two separate genetic perturbation screens: one involving RPE1 cells with 1,543 perturbations and another with K562 cells, comprising 1,092 perturbations. Each dataset contained data from over 160,000 cells [26]. The performance of AttentionPert is compared against baseline models, as detailed in Table 2. The results demonstrate that AttentionPert outperforms all baselines in all metrics.

**Table 2:** Seen 0/1 comparison of 3 metrics on RPE1 and K562 datasets.

| Data | Model | MSE(DE) | rel-MSE(DE) | $\rho_{\Delta}(\text{all})$ |
| --- | --- | --- | --- | --- |
| RPE1 | Ctrl | 0.290 | 100 | -0.017 |
| | CPA | 0.281 $\pm 0.001$ | 97.0 $\pm 0.4$ | 0.109 $\pm 0.025$ |
| | GEARS | 0.143 $\pm 0.005$ | 49.3 $\pm 1.6$ | 0.527 $\pm 0.025$ |
| | Ours | **0.132** $\pm 0.005$ | **45.7** $\pm 1.7$ | **0.532** $\pm 0.010$ |
| K562 | Ctrl | 0.211 | 100 | -0.014 |
| | CPA | 0.205 $\pm 0.000$ | 97.3 $\pm 0.1$ | 0.088 $\pm 0.015$ |
| | GEARS | 0.137 $\pm 0.003$ | 65.1 $\pm 1.2$ | 0.327 $\pm 0.015$ |
| | Ours | **0.132** $\pm 0.003$ | **62.4** $\pm 1.3$ | **0.340** $\pm 0.012$ |

Results in Supplementary Section E.1 show that AttentionPert beats all the baselines on all other 4 splits of each dataset, which underscores the robustness and adaptability of AttentionPert under diverse test conditions.

### 3.3 Discovering Genetic Interactions

Non-Gaussianity in residual errors reveals outlier complexity in data not captured by the model. Here, we analyze AttentionPert’s residual errors to diagnose any failure cases in the model, while also revealing nuanced genetic interactions, outlier complexity in the regulation of some genes, and gene modules with correlated errors (Figure 5). AttentionPert achieves near-zero error on genes GMFG, ACP5, POU3F2, and NGRN, regardless of perturbation condition, while the genes RUNX1T1, MS4A3, LGI2, and CTSL display high error across all conditions, indicating nuanced regulatory behavior unrelated to gene perturbations. Surprisingly, perturbations to transcription factor families such as the CEB* family and FOX* family have much lower residual error across all genes, suggesting that these are degenerate cases where the complexity of the transcriptome decreases in their absence. In contrast, the two perturbation cases with BCL2L11 – a signaling protein involved in apoptosis regulation – have much higher residuals than average, indicating an induced incoherent system or cellular decay.

**Figure 5.**
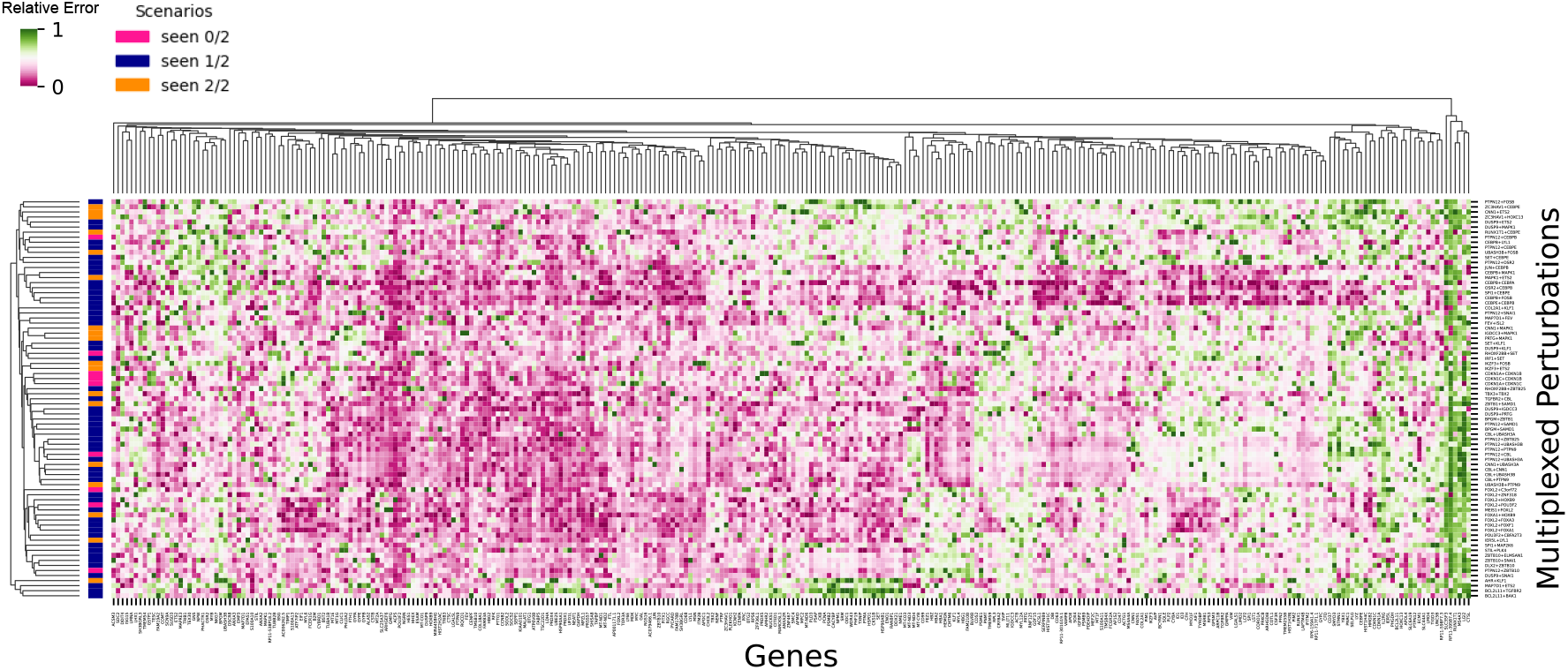
Gene-specific residual errors of AttentionPert. The results are shown over tested multiplexed perturbation conditions in the Norman dataset and the union set of the top 20 DE genes of these perturbations. Scenarios of out-of-domain generalization are plotted on the left. Residual errors are log-transformed and column-scaled.

Notably, the residuals are uncorrelated with the perturbation domain splits (Figure 5), and uncorrelated with the structure of the Gene Ontology graph (Supplementary Figure 11). Complementing AttentionPert’s SOTA performance on all domain generalization tasks (Table 1), this gives us high confidence that AttentionPert generalizes extremely well, and that high error genes and perturbations are indicative of complex and nuanced genetic mechanisms.

Furthermore, non-additive effects arising from the complex interactions between multiple perturbed genes underscore the necessity for a model that can accurately and integrally handle multigene perturbations. Figure 6 reveals the ability of AttentionPert to capture the non-additive nature of multiplexed genetic perturbations. When predicting a seen 2/2 perturbation: FOXA1 and HOXB9, AttentionPert is adept at learning the combined response of these genes. This approach differs markedly from a simple additive model, which merely sums the transcriptional effects of each gene perturbed independently.

**Figure 6.**
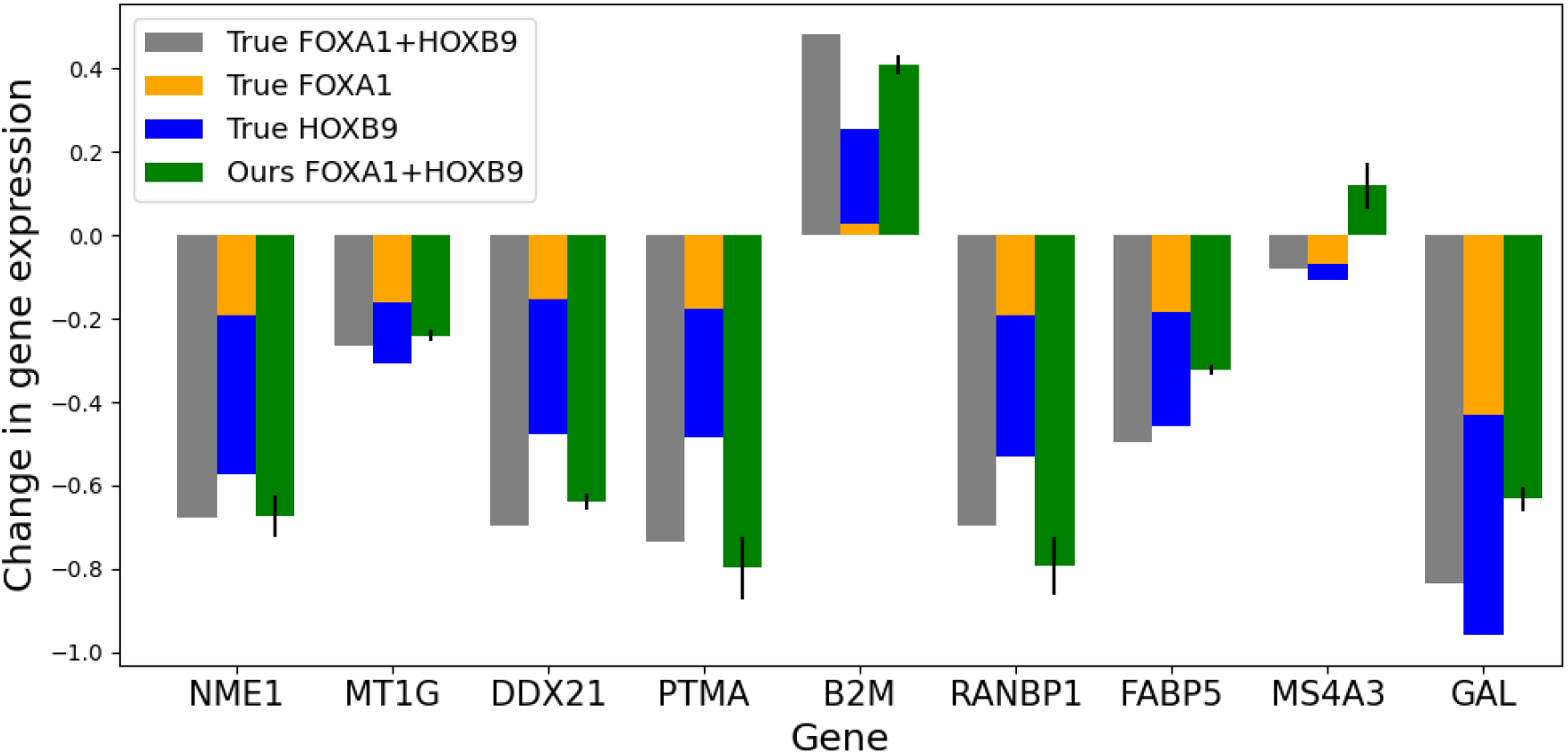
Change in gene expression after perturbing the combination FOXA1 and HOXB9. The gray bars indicate the true mean post-perturbation gene expression change. The orange and blue bars show the sum of true shifts for the two individual single-gene perturbations. The green bars show the prediction made by AttentionPert.

### 3.4 Ablation study

The primary distinction between AttentionPert and the SOTA GEARS lies in the expression shift function, **g**_*θ*_. Besides the model architecture, AttentionPert differs from GEARS in three aspects: first, we discard the cross-gene layer utilized by GEARS; second, we match the number of potentially perturbed genes to the size of the genes in the dataset, while GEARS includes a different set of potential perturbed genes, adding numerous nodes to the GO graph that are not perturbed in actual datasets; thirdly, we utilize pre-trained Gene2Vec embeddings for initializing all gene embeddings in AttentionPert, while GEARS employs random initialization. The ablation study presented in Table 3 demonstrates that AttentionPert achieves the lowest MSE(DE) in seen 0/2 scenario, which indicates that the enhanced performance of AttentionPert is not solely attributable to any one of these three changes or their combination. Therefore, the design of AttentionPert is essential for its improved performance.

**Table 3:** Ablation experimental results comparing other differences from GEARS. *CG* means utilizing the cross-gene decoding layer. *Rd* means reducing the potentially perturbed genes. *G2V* is initializing the gene embeddings with Gene2Vec. The original GEARS method utilizes cross-gene without reducing genes or using Gene2Vec.

| Model | CG | Rd | G2V | Seen 0/2<br>MSE(DE) | Seen 1/2<br>MSE(DE) | Seen 2/2<br>MSE(DE) |
| --- | --- | --- | --- | --- | --- | --- |
| GEARS | ✓ | | | 0.219 ±0.011 | 0.221 ±0.007 | 0.130 ±0.007 |
| GEARS | ✓ | | ✓ | 0.293 ±0.010 | 0.449 ±0.009 | 0.391 ±0.006 |
| GEARS | ✓ | ✓ | | 0.235 ±0.006 | 0.234 ±0.008 | 0.117 ±0.004 |
| GEARS | ✓ | ✓ | ✓ | 0.194 ±0.012 | 0.225 ±0.007 | 0.110 ±0.003 |
| GEARS | | ✓ | | 0.246 ±0.022 | 0.215 ±0.011 | **0.087** ±0.003 |
| GEARS | | ✓ | ✓ | 0.208 ±0.015 | 0.211 ±0.005 | 0.094 ±0.004 |
| Ours | | ✓ | ✓ | **0.159** ±0.012 | **0.193** ±0.003 | 0.097 ±0.008 |

In our study, we conduct ablation experiments on the two encoders: PertWeight and PertLocal. The results presented in Table 4 reveal that while each encoder contributes to the model’s performance, neither encoder individually outperforms the effectiveness of the integrated model that combines both encoders. This underscores the synergistic effect of incorporating both the PertWeight and PertLocal encoders into AttentionPert.

**Table 4:** Ablation experimental results on AttentionPert. We compare our model with/without each of the three components: PertLocal (*PL*), PertWeight (*PW*), and Gene2Vec (*G2V*) initialization.

| Model | PL | PW | G2V | Seen 0/2<br>MSE(DE) | Seen 1/2<br>MSE(DE) | Seen 2/2<br>MSE(DE) |
| --- | --- | --- | --- | --- | --- | --- |
| only-PL | ✓ | | ✓ | 0.246 ±0.002 | 0.388 ±0.004 | 0.324 ±0.002 |
| only-PW | | ✓ | ✓ | 0.172 ±0.009 | 0.202 ±0.004 | 0.120 ±0.007 |
| Ours | ✓ | ✓ | | 0.189 ±0.026 | 0.203 ±0.009 | 0.105 ±0.018 |
| Ours | ✓ | ✓ | ✓ | **0.159** ±0.012 | **0.193** ±0.003 | **0.097** ±0.008 |

Besides, we conduct experiments where AttentionPert does not use Gene2Vec. Results in Table 4 and Table 3 show that it beats GEARS without using Gene2Vec but has worse results than the final version of AttentionPert where we use Gene2Vec to initialize gene embeddings.

Another interesting finding is that the original version of GEARS using Gene2Vec performs worse, while the reduced version of GEARS using Gene2Vec archives a better performance as shown in Table 3. This is probably because the number of possible perturbed genes used by the original GEARS is much larger than the actual number of genes in the Norman dataset. A pre-trained gene context representation undermines its ability when dealing with a smaller set of genes.

Ablation study results for other splits are detailed in Supplementary Section E.2.

## 4 Conclusion

We have proposed a comprehensive model AttentionPert that effectively integrates attention mechanisms to predict the unique effects of single or multiple perturbed genes. By incorporating the PertWeight and PertLocal encoders that complement each other with encodings representing global and local perturbational effects respectively, we have successfully addressed the challenges of predicting transcriptional responses to genetic perturbations. AttentionPert not only outperforms existing state-of-the-art methods in terms of accuracy and handling diverse gene perturbations but also shows exceptional capability in predicting out-of-distribution scenarios where all simultaneously perturbed genes are unseen. The demonstrated superior performance across multiple datasets highlights its potential in facilitating medical research and treatment development. Besides, detailed gene- and perturbation-specific experimental analyses provide valuable insights into the complex genetic interactions inherent in multi-gene perturbations, which clarify the intricate dynamics of gene interplay under various perturbation conditions. We anticipate further exploration and application of the computational method to predict cellular responses of novel genetic perturbations in personalized medicine and genetic research.

## A Method Details

### A.1 Data preprocessing and loss functions

We use the three perturbational effects datasets [22, 26] preprocessed by GEARS [27] to evaluate our model. The preprocess of GEARS first normalizes the expression values, discard low-varying genes, then samples an unperturbed gene expression 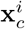 from the set *{***x**^*t*^, *t* = 1, …, *n*_control_*}* to be 1-to-1 correspondent with each post-perturbation expression vector 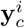 where *i* = 1, …, *n*_*c*_. For the Gene Ontology (GO), we use the GO datasets provided by the SOTA method and also the Gene Ontology knowledgebase [4, 31, 30]. After aligning the genes, we have successfully found more GO terms for those genes not shown in the GO graph provided by GEARS.

We also utilize Gene2Vec [9] which are 200-dimension vector representations for human genes. We use a Gaussian distribution to sample the vector for genes shown in each of the datasets that cannot be aligned with the gene symbols provided by Gene2Vec. Formally, assuming the gene vector is **v**_*g*_ where *g* ∈ *𝒢*is theset of gene symbols of Gene2Vec, for any unknown gene *x* ∈*/ 𝒢*, its representation is sampled as:

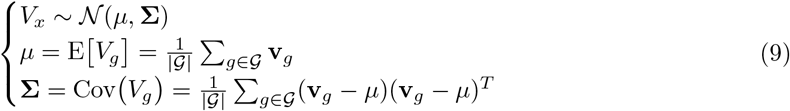

We follow the loss functions used by GEARS [27]. During the training, GEARS uses a loss

function that measures the distance between each predicted expression 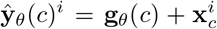 and the ground-truth 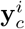, which is:

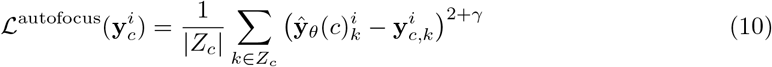

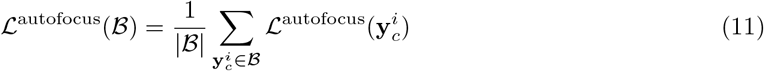

where ℬ is a batch of post-perturbation expressions in the training process, *Z*_*c*_ is the set of genes with non-zero expressions in at least 1 cell under the perturbation condition *c*, and *γ* is a hyper-parameter which = 2 in GEARS experiment setup.

GEARS also combines a direction-aware loss with a parameter *λ* = 0.1 with the autofocus loss:

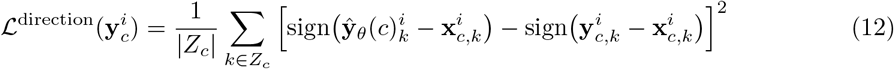

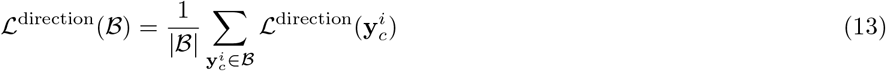

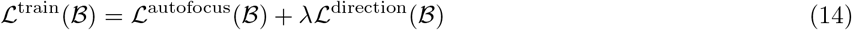

In addition, the log variance in GEARS is taught to act as a proxy for model uncertainty by promoting its increase when errors are significant. This is achieved through a gene-specific layer that predicts the log variance for each gene, which is incorporated into the learning process via a modified Bayesian neural network loss, as detailed in [18]. To estimate the post-perturbation gene expression values, a Gaussian likelihood approach is employed. The uncertainty score serves as an indicator of the confidence level of the model when predicting outcomes for new perturbations.

### A.2 An intuitive geometric illustration of our method

Our model, AttentionPert, operates on the fundamental assumption that each gene’s unperturbed state can be represented as a vector within a *D*-dimensional hidden space. When certain genes undergo perturbation, we map these perturbation conditions to this high-dimensional space. This mapping alters the basal states of the genes into perturbed states, which are then decoded into scalar values representing the changes in post-perturbation expressions.

AttentionPert comprises two principal encoders: PertLocal and PertWeight. PertLocal is responsible for locally perturbing the basal states of genes. In contrast, PertWeight is designed to calculate both general and gene-specific effects on a more global scale. The terms *locality* and *globality* in this context are defined based on a gene-gene relationship graph. For our model, this graph is the Gene Ontology (GO) graph, which may be used in its standard form or in an augmented version. This entire sequence of events in the hidden space is illustrated in Fig 7.

**Figure 7.**
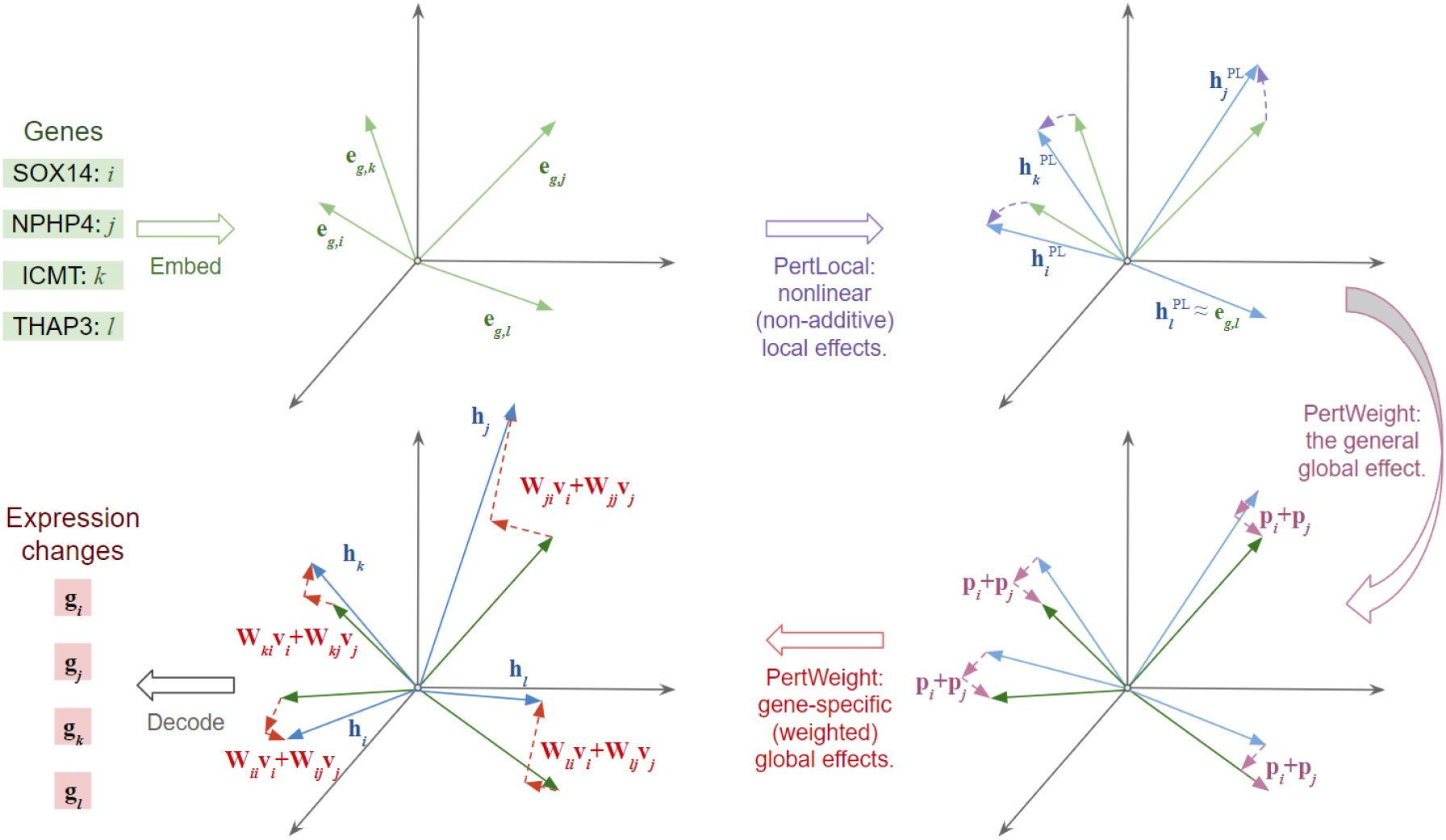
The geometric illustration of AttentionPert. First, genes *i, j, k, l* are embedded into a high-dimensional hidden space as vectors **e**_*g,i*_, **e**_*g,j*_, **e**_*g,k*_, **e**_*g,l*_. Assume that genes *i* and *j* are perturbed and are both local to *k* but distant from *l* in the GO graph. Then PertLocal locally (with non-additivity) perturbs each gene-representing vector into **H**^PL^. After that, PertWeight adds the general global effect **p**_*i*_ + **p**_*j*_ to all vectors and finally adds a gene-specific global effect to each vector, resulting the encoding **H** = **H**^PL^ + **H**^PW^. The encoding for each gene is subsequently decoded into a scalar representing the expression change.

### A.3 Graphs and GNNs

The GO graph is constructed by incorporating the shared GO terms of each pair gene [27], and those genes in the dataset that have no GO terms or have no intersected ones with other genes are still added to the graph as outlier nodes. Formally, for each pair of genes *i* and *j* ∈ *{*1, 2, …, *K}*, denote the sets of corresponding Gene Ontology terms to be *𝒩*_*i*_ and *𝒩N*_*j*_. Then the edge weight between these 2 genes is defined as **J**_*ij*_ = |*𝒩*_*i*_ ∩ *𝒩*_*j*_|*/*|*𝒩*_*i*_ ∪ *𝒩*_*j*_|, which is the fraction of shared Gene Ontology terms between the 2 genes. After that, for each gene *i*, the *H*_pert_ = 20 gene *j*’s with the highest **J**_*i,j*_ are selected to construct the Gene Ontology graph, and this selection gives each edge a direction since the top-weighted neighboring is not bidirectional. Hence the graph 𝒥 is an edge-weighted directed graph that uses the Gene Ontology to construct gene-gene interactions, as shown in Figure 2.

Unlike GEARS, we also reduce the GO graph by not including possible perturbed genes that are not actually perturbed in the datasets. Instead of using all possible perturbed genes, we include all the genes in each of the three evaluation datasets to create the GO graph. In conclusion, the GO graph used by GEARS is the same graph of all possible perturbed genes for all three datasets, while our reduced GO graph is unique for each dataset that only contains all the genes from it. Figure 10 shows that the GEARS using the reduced version of the GO graph usually makes better results than its original version.

We use SGConv [19] as the four GNN encoders in our model. A *t*-hop SGConv with a learnable weight matrix 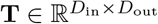 outputs:

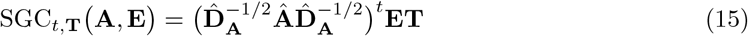

where **Â** = **A** + **I** denotes the adjacency matrix with inserted self-loops and 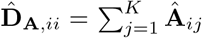 its diagonal degree matrix. The output is in 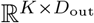.

### A.4 Non-additive features

As a perturbational effects prediction method, AttentionPert has theoretical non-additive features. Non-additivity requires the predicted *g*_*θ*_(*c*)_*k*_ not always equal to 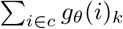 for any multi-gene perturbation *c*, and the non-linear coeffect of multiple perturbed genes can be learned.

The PertWeight encoder generates an additive encoding of any multi-gene condition *c*, formally,

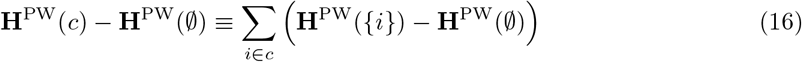

Note that here **H**^PW^(∅) ≡ 0. However, PertLocal encoder utilizes the non-additive bias layer to avoid a linear combination of perturbed genes,

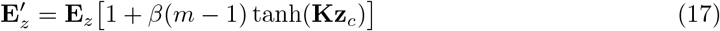

where gene indices *i, j* ∈ *{*1, …, *K }*. Since the function tanh is non-linear and *m* is the number of perturbed genes:

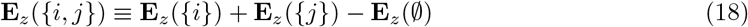

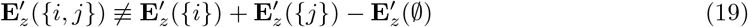

Therefore the PertLocal encoding is non-additive across all perturbed genes in any given condition *c*:

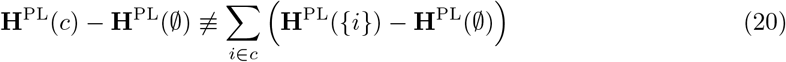

This implies that the final output is not additive for multi-gene perturbations either.

## B Evaluation Metric Details

The primary metric for evaluating the performance under each perturbation condition *c* is the Mean Squared Error (MSE). This metric measures the deviation between the expected prediction, approximated as the average predicted expressions E [*Y*_*θ*_(*c*)], and the actual ground-truth post-perturbation expression E[*Y*_*c*_] focusing on the top 20 differentially expressed (DE) genes, denoted as 𝒟_*c*_ 1, …, *K* where |𝒟_*c*_| = 20. The expected expression for both predicted and actual samples is calculated as the mean value across all cells under the given perturbation condition *c*. Formally:

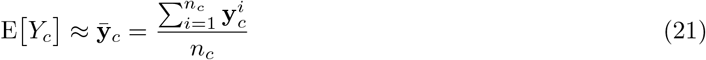

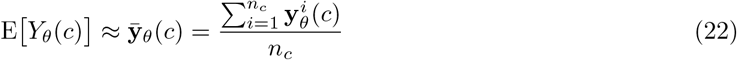

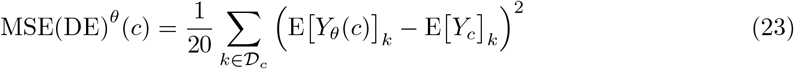

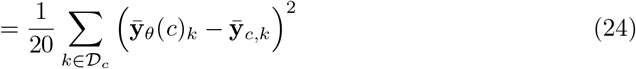

Due to the substantial cost associated with CRISPR gene-perturbation screens, there is a pressing need for an in-silico method capable of predicting transcriptomic outcomes of novel single or multi-gene perturbations absent from existing datasets. In our experimental design, cells under a specific perturbation condition *c* are entirely allocated to either the training and validation set or the test set. This allocation strategy effectively simulates the challenge of predicting perturbations that the model has not previously encountered. For any given dataset, we define two distinct sets of perturbation conditions: 𝒞_tv_, representing the training and validation set, and 𝒞_test_, representing the test set. As all unperturbed cells are considered known and are therefore included in the training and validation set, the control condition is defined as *c*_control_ = ∅ and is part of 𝒞_tv_. This approach ensures a clear demarcation between training/validation and testing scenarios, crucial for evaluating the model’s efficacy in handling novel perturbations.

When the number of perturbed genes is only 1, there is only one testing scenario that is called *seen 0/1*, because any perturbation condition containing only 1 gene which either shows in the train and validation set or shows in the test set. We denote such a scenario as 𝒮_0*/*1_ = *{c* ∈ *𝒞*_test_||*c*| = 1*}*.

When considering the combination of multiple perturbed genes, however, there could be multiple testing scenarios, depending on the experimental exposure of the genes to perturbations during training. For *s*-gene perturbations test where *s >* 1 (in real-life datasets we only have *s* = 2), there could be *s* + 1 scenarios, which is 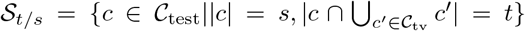 where *t* = 0, 1, …*s* meaning the number of genes in this perturbation condition that has shown in the train and validation sets. By that definition, over a *s*-perturbation test, a scenario with smaller *t* is a more OOD case and harder for the model to predict. The top 20 DE MSE metric of a test scenario 𝒮_*t/s*_ can be formulated as:

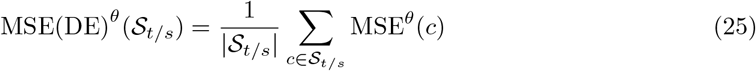

The relative MSE (rel-MSE) in percentage, which is the proportion relative to the Ctrl model, is calculated by:

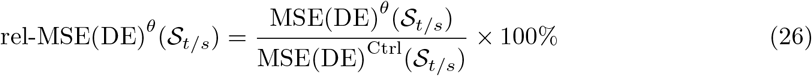

and by definition of the Ctrl model that only outputs the average control sample:

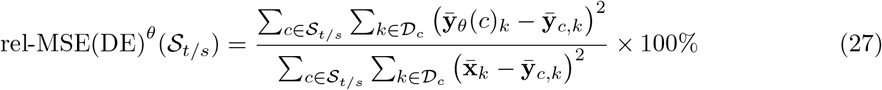

In addition to MSE, we assess our model’s performance using the Pearson correlation coefficient, which measures the relationship between the predicted and actual post-perturbation expression shifts. Formally:

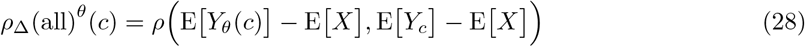

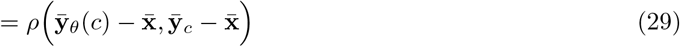

where *ρ*(*·, ·*) is the Pearson correlation coefficient of 2 vectors. The metric for a set of perturbations is also the average *ρ*_Δ_(all)^*θ*^ over all perturbations in the given set. This coefficient is calculated across all genes to gauge the model’s proficiency in predicting overall expression changes.

Beyond MSE and the Pearson score, we employ two additional metrics for a more comprehensive evaluation of predictive error rates of top 20 DE genes. The proportion of top 20 DE genes for a given perturbation predicted to change in an opposite direction of the ground truth is formally:

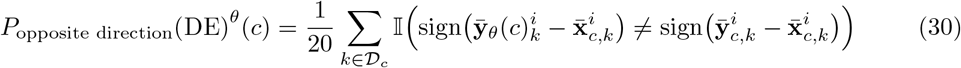

The proportion of DE genes for which the predicted post-perturbation gene expression falls beyond one standard deviation of the actual post-perturbation expression distribution mean is formally:

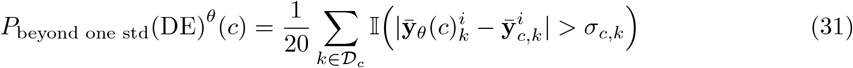

Here 𝕀(*·*) is the indicator function which equals 1 if the variable is true and 0 otherwise. The standard deviation of perturbation *c* on gene *k*: *σ*_*c,k*_ is over the set of ground truth expressions : 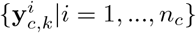. Both metrics are averaged as the score of a set of perturbations as well.

## C Hyperparameter Search

We tune and fix the best set of hyperparameters on the validation set of the first split of the Norman dataset [22]. We search hyperparameters in ranges: the number of hops of SGConvs *t* ∈ *{*1, 2*}*; the network hidden size *D* ∈ *{*32, 64, 128*}*; for the multi-head attention, the number of heads *H* ∈ *{*1, 2, 4, 8*}* and the dimension of each head *d*_*q*_ ∈ *{*8, 16, 32, 64, 128, 256*}*, where *H × d*_*q*_ ∈ [64, 256]; the minimum edge weight parameter in the augmented GO graph *α* ∈ *{*0.25, 0.5, 0.75, 1.0*}*; the NA-bias parameter *β* ∈ *{*0.001, 0.005, 0.01, 0.05*}*; batch size in *{*32, 64, 128, 256, 512*}*. The best hyperparameters are listed in the main body, where *t* = 1, *D* = 64, *H* = 2, *d*_*q*_ = 64, *α* = 0.75, *β* = 0.05. To ensure a fair comparison with the GEARS model, we optimized the batch size, selecting 128 as the most effective value for both models. Additionally, we search the number of top-weighted edges retained in the Gene Ontology (GO) graph. The results indicated similar performance across a range of values 10, 15, 20, 25, leading us to select 20, aligning with the configuration used in GEARS. Our model is versatile in its representation of prior knowledge regarding gene-gene relationships. While it can utilize various types of networks, such as protein-protein interactions or gene coessentiality networks, we choose to use the GO graph, as detailed in Supplementary Note 2 by GEARS [27]. This decision is driven by the desire to maximize gene coverage and maintain consistency for a fair comparison with GEARS.

## D Datasets details

Table 5 shows the details of each dataset, including the numbers of different perturbations, corresponding cell samples, and genes in each dataset covered in the Gene Ontology database or Gene2Vec. During the experiments, each dataset has 5 different splits for training, validation, and testing. We use the same split ratio of genes that the SOTA method conducts its experiments, where for both single-gene and multi-gene cases, the number of perturbed genes in the train and validation set is 75% of the total number of perturbed genes of the dataset. This makes 5 different sets of numbers of 3 scenarios in the 2-gene perturbational dataset [22] but the same number of the only 1 scenario for 1-gene datasets [26], shown in Table 6.

**Table 5:**
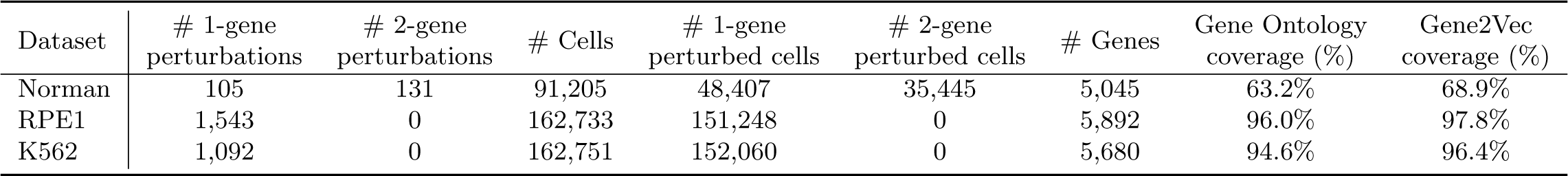
Details of the perturbational effects datasets. Here # means the number of something. The Norman dataset is a two-gene perturbation dataset [22]. The RPE1 and K562 datasets are single-gene perturbation datasets [26]. All three datasets are pre-processed by GEARS [27].

**Table 6:**
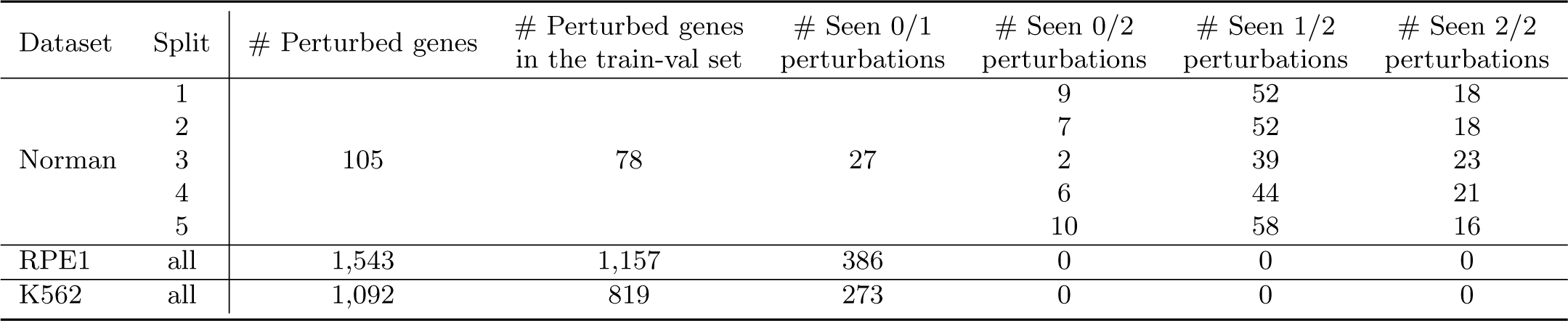
Different splits of Norman, RPE1 and K562 datasets. Here # means the number of something. For each dataset, the gene ratio of the train-validation set is 75%. The Norman dataset is a two-gene perturbation dataset [22] that contains 3 scenarios in the test set. The RPE1 and K562 datasets are single-gene perturbation datasets [26] with only seen 0/1 scenario. GEARS pre-processes all three datasets [27].

| Dataset | Split | # Perturbed genes | # Perturbed genes in the train-val set | # Seen 0/1 perturbations | # Seen 0/2 perturbations | # Seen 1/2 perturbations | # Seen 2/2 perturbations |
| --- | --- | --- | --- | --- | --- | --- | --- |
| Norman | 1 | 105 | 78 | 27 | 9 | 52 | 18 |
|  | 2 |  |  |  | 7 | 52 | 18 |
|  | 3 |  |  |  | 2 | 39 | 23 |
|  | 4 |  |  |  | 6 | 44 | 21 |
|  | 5 |  |  |  | 10 | 58 | 16 |
| RPE1 | all | 1,543 | 1,157 | 386 | 0 | 0 | 0 |
| K562 | all | 1,092 | 819 | 273 | 0 | 0 | 0 |

## E Additional Experiment Results

### E.1 Detailed results for comparison on all 5 splits of 3 datasets

The difficulty across different splits of the same dataset can vary considerably since we do not randomly split each dataset across all cells. Instead, we ensure that all perturbations in the test set are not encountered during the training phase, leading to significant variability in model performance across different experimental splits. Therefore, rather than averaging evaluation metrics across all splits, we present a split-by-split comparison of the three datasets. Table 7 shows the detailed MSE and *ρ*_Δ_ results on the other 4 splits of the Norman dataset (results for the first split are shown in the main body). Besides, percentages of DE genes with an opposite direction and beyond one standard deviation results of all 5 splits of the Norman dataset are shown in Fig 8. Results show that AttentionPert outperforms other methods in most of the metrics on every split when dealing with multi-gene perturbations.

**Table 7:**
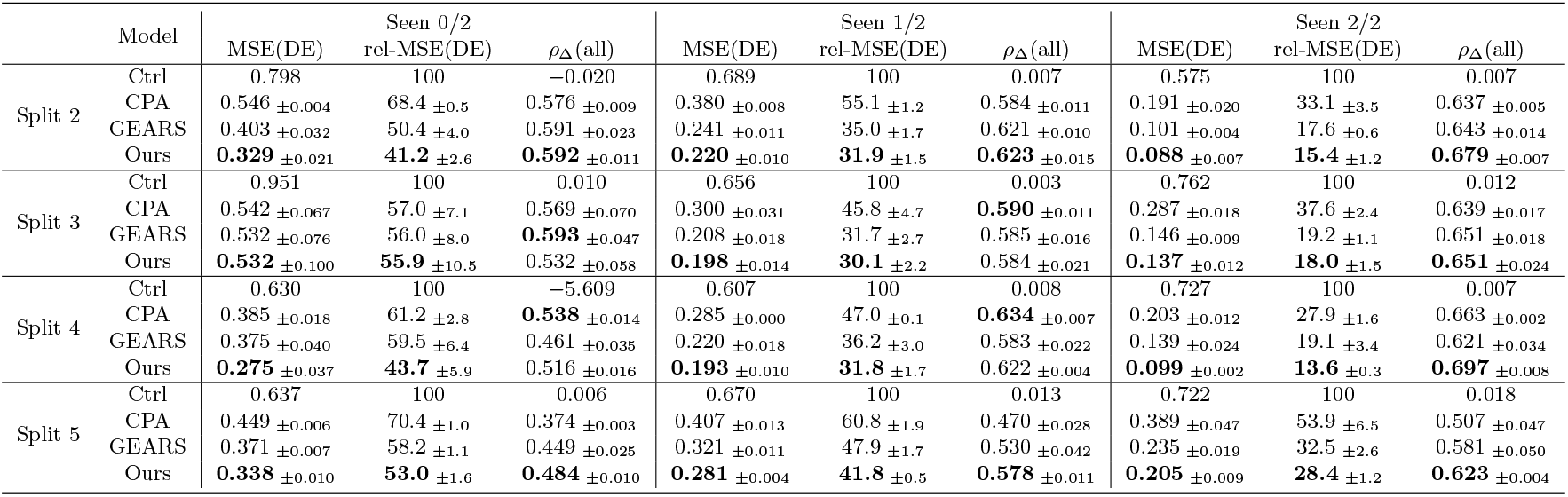
Seen 0/2, seen 1/2, and seen 2/2 comparison of MSE, relative MSE (%) and *ρ*_Δ_ on other 4 splits of Norman dataset. MSE stands for the MSE of the top 20 DE genes. The relative mean squared error (rel-MSE) is in percentage (%). The *ρ*_Δ_ score refers to the Pearson score between the predicted shift and the ground-truth shift on all genes. We record the mean and the standard deviation over 5 independent experiments. The best performances of each split are marked in bold.

The detailed MSE, rel-MSE and *ρ*_Δ_ results on the other 4 splits of RPE1 and K562 datasets (results for the first split are shown in the main body) are shown in Table 8, while error-rate representing results of all 5 splits of these two datasets are shown in Fig 9. Based on the experimental performances, AttentionPert also beats other methods in most of the metrics on every split for single-gene perturbation datasets.

**Table 8:**
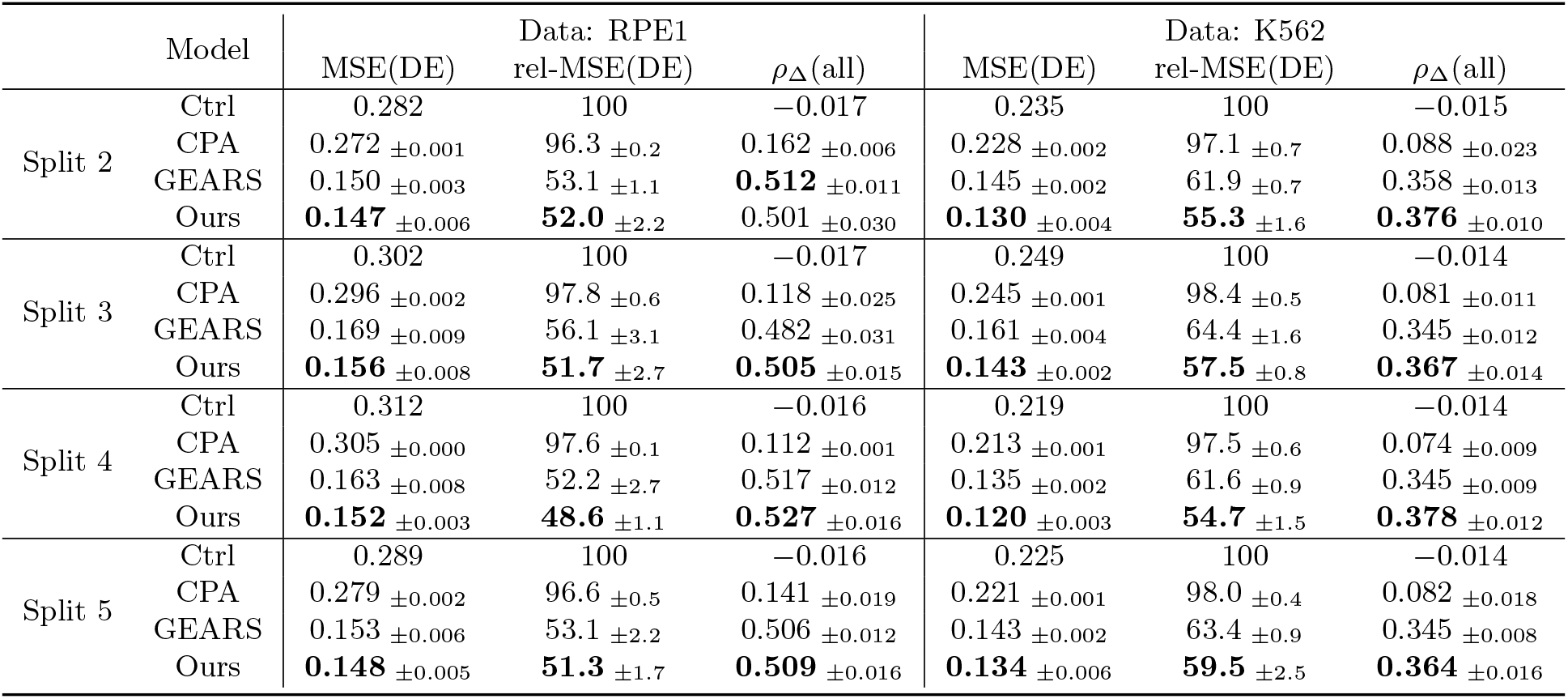
Seen 0/1 MSE, relative MSE (%) and *ρ*_Δ_ results on other 4 splits of RPE1 and K562 datasets. The best performances of each split are marked in bold.

### E.2 Detailed results of the ablation study

The rel-MSE results for seen 0/2, seen 1/2 and seen 2/2 scenarios of the ablation experiments are shown in Figure 10. We can see that AttentionPert does not beat all other revised GEARS models in seen 1/2 and seen 2/2 metrics. However, the results for seen 0/2 demonstrate that for the most OOD case where either of the perturbed genes is unseen when training, our model outperforms the revised version of GEARS and also any other methods. Therefore we still conclude that our model generally performs better than any revised version of GEARS.

**Figure 8.**
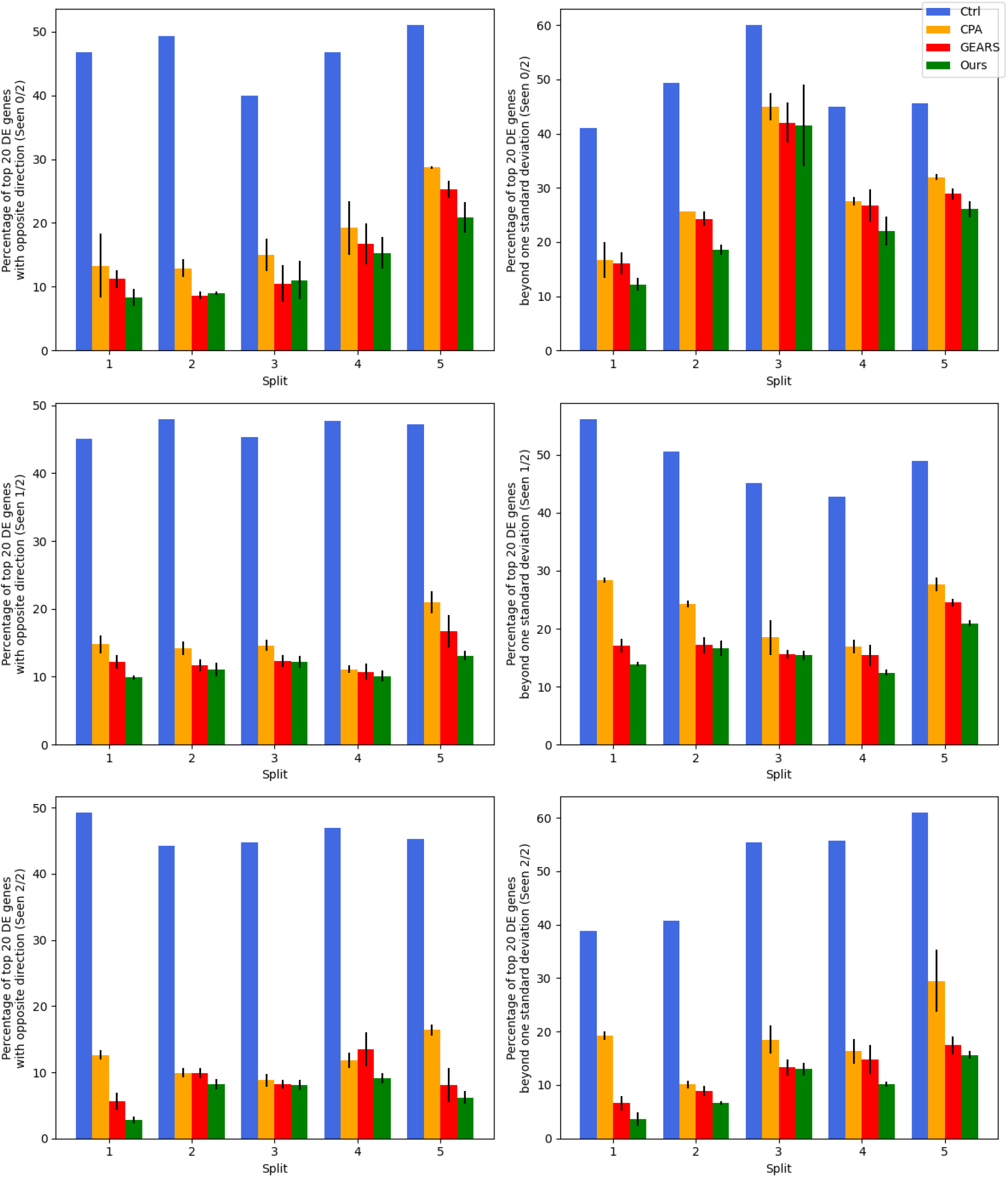
Comparison of two error-rate representing metrics on Norman dataset.

**Figure 9.**
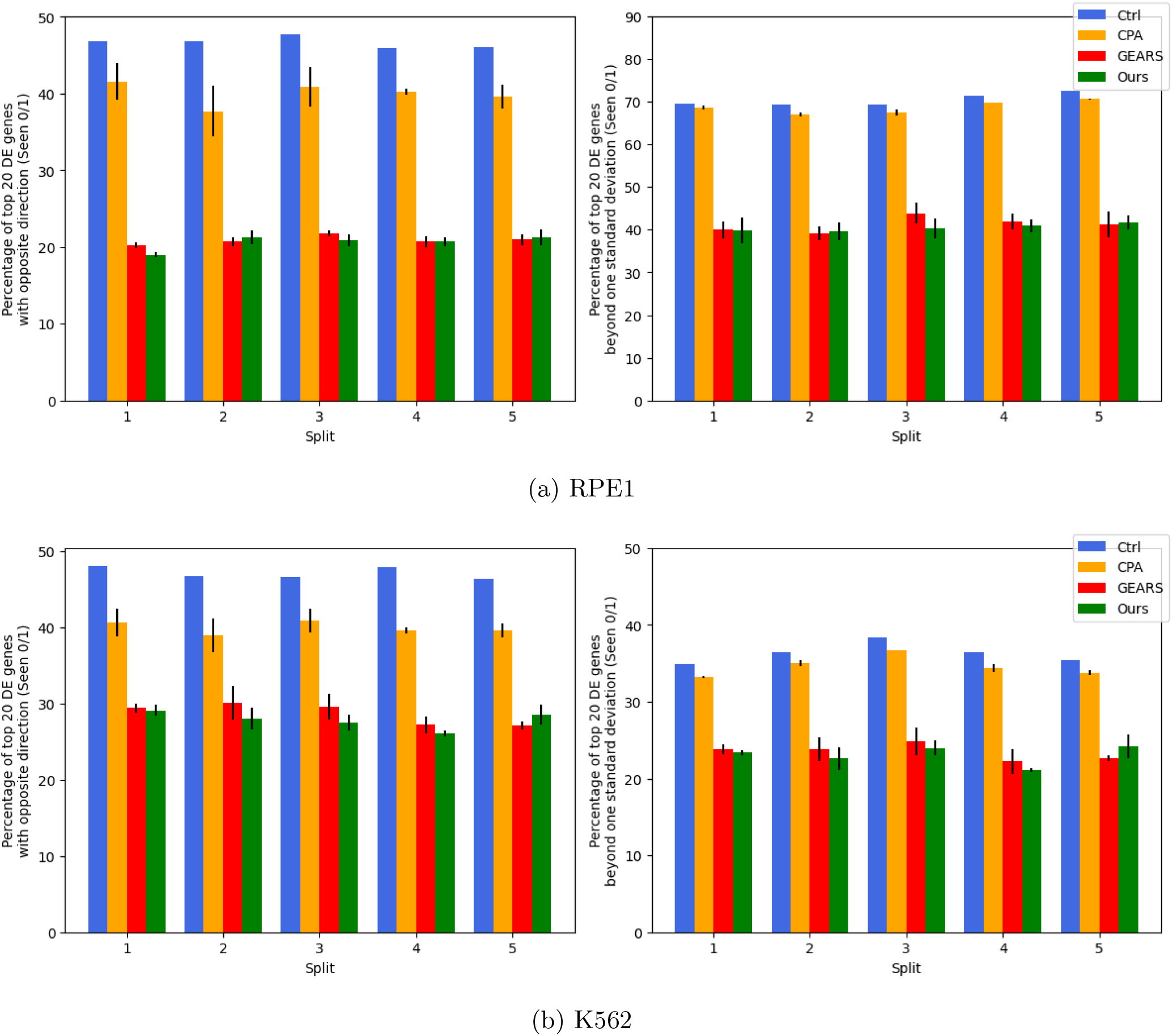
Comparison of two error-rate representing metrics of RPE1 (a) and K562 (b) datasets. Both are single-gene perturbation datasets with only seen 0/1 to compare.

**Figure 10.**
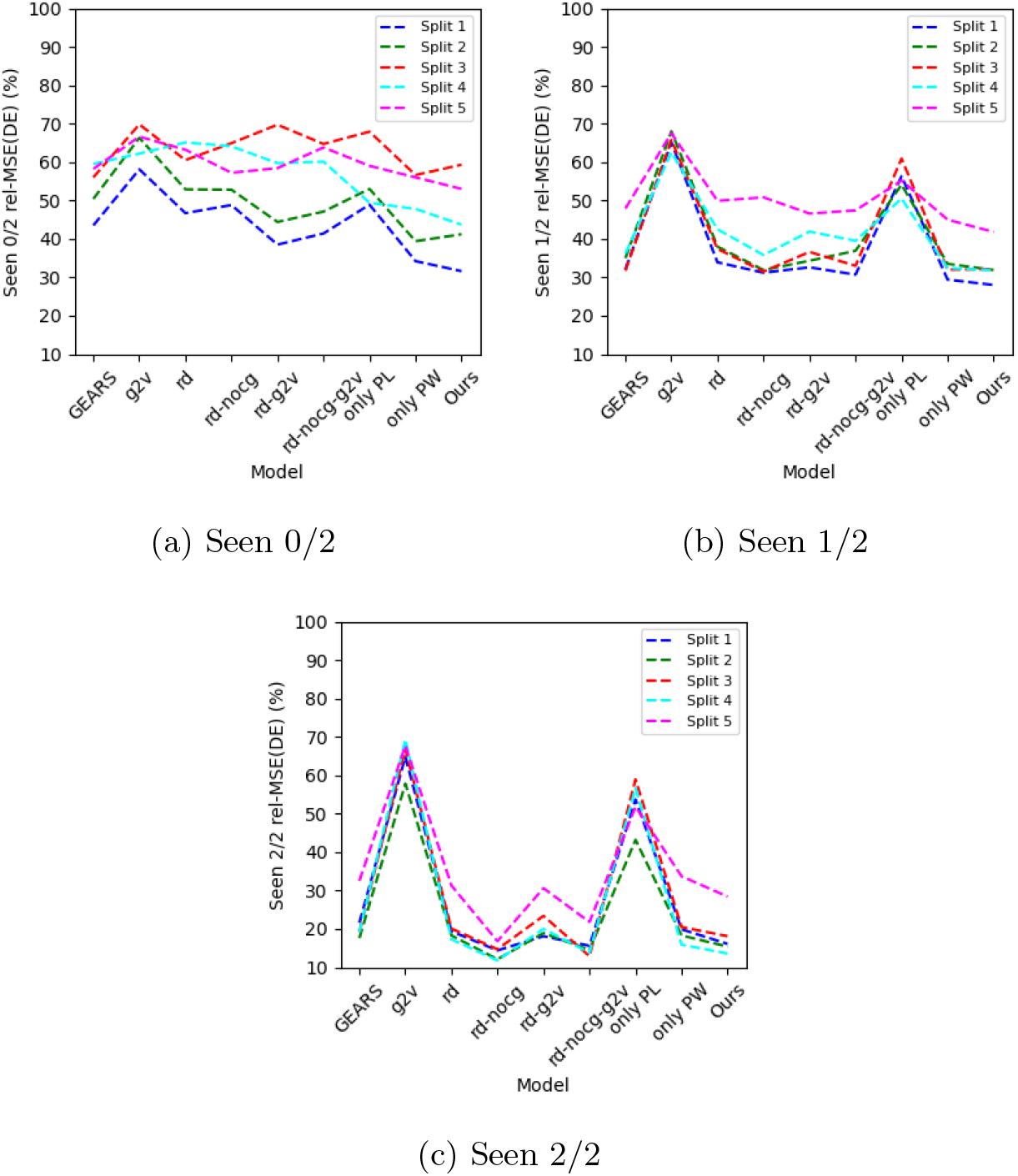
The ablation study comparing rel-MSE (%) of top 20 DE genes over seen 0/2 (a), seen 1/2 (b) and seen 2/2 scenarios (b), on 5 splits of Norman dataset. Here, *g2v* is GEARS that also uses Gene2Vec to initialize the gene embedding, *rd* model is GEARS changed by merely reducing the possible perturbed genes to the genes in the dataset, and *nocg* is GEARS without the cross-gene layer. For our design, *only PL/PW* means the model with only the PertLocal/PertWeight encoder.

**Figure 11.**
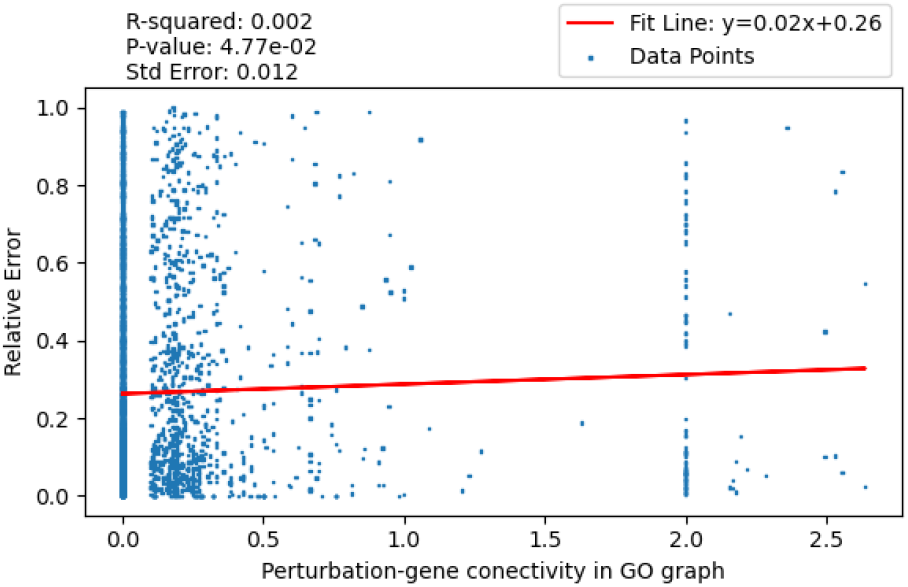
Relative error of AttentionPert versus connectivity of perturbation-gene pair. The connectivity is defined as the sum of edge weights between the perturbed genes and the affected gene in the Gene Ontology graph.

